# Identification of HPV oncogene and host cell differentiation associated cellular heterogeneity in cervical cancer via single-cell transcriptomic analysis

**DOI:** 10.1101/2023.08.10.552878

**Authors:** Yingjie Li, Cankun Wang, Anjun Ma, Abdul Qawee Rani, Mingjue Luo, Jenny Li, Xuefeng Liu, Qin Ma

## Abstract

Human Papillomaviruses (HPVs) are associated with around 5-10% of human cancer, notably nearly 99% of cervical cancer. The mechanisms HPV interacts with stratified epithelium (differentiated layers) during the viral life cycle, and oncogenesis remain unclear. In this study, we used single-cell transcriptome analysis to study viral gene and host cell differentiation-associated heterogeneity of HPV-positive cervical cancer tissue. We examined the HPV16 genes - E1, E6, and E7, and found they expressed differently across nine epithelial clusters. We found that three epithelial clusters had the highest proportion of HPV-positive cells (33.6%, 37.5%, and 32.4%, respectively), while two exhibited the lowest proportions (7.21% and 5.63%, respectively). Notably, the cluster with the most HPV-positive cells deviated significantly from normal epithelial layer markers, exhibiting functional heterogeneity and altered epithelial structuring, indicating that significant molecular heterogeneity existed in cancer tissues and that these cells exhibited unique/different gene signatures compared with normal epithelial cells. These HPV-positive cells, compared to HPV-negative, showed different gene expressions related to the extracellular matrix, cell adhesion, proliferation, and apoptosis. Further, the viral oncogenes E6 and E7 appeared to modify epithelial function via distinct pathways, thus contributing to cervical cancer progression. We investigated the HPV and host transcripts from a novel viewpoint focusing on layer heterogeneity. Our results indicated varied HPV expression across epithelial clusters and epithelial heterogeneity associated with viral oncogenes, contributing biological insights to this critical field of study.

## Background

Human Papillomavirus (HPV) infection causes the most common sexually transmitted diseases and up to 5% of human cancer, including anogenital cancer, and head & neck cancer ^1–3^. Specifically, persistent HPV infection in the host epithelium is one of the critical steps of cervical cancer development ^4,5^. Current prophylactic HPV vaccines have enabled the prevention of initial infection and HPV-associated malignancies. However, HPV vaccination rates vary by state in the US (from 31% to 79% of adolescents being HPV UTD (up-to-date)), certain high-risk HPV types are not covered in vaccines, and these vaccines will not eliminate viruses from infected individuals. There is no available effective treatment for HPV persistent infection, due to the precise processes and underlying mechanism of the natural HPV infection largely unknown ^6^. Thus, HPV-associated human diseases and cancers remain a big challenge in the future 20-40 years ^7–10^.

The HPV life cycle is highly regulated in the host-stratified cutaneous and mucosal epithelia ^11–13^. In the cervix, four cell layers are involved in the epithelium, which are the basal layer, parabasal layer, squamous cells layer, and matured squamous layer. Persistent HPV infectious cycle requires maintenance of HPV genome DNA as low copy number extrachromosomal form in the dividing basal cells and amplifying progeny viral genomes in differentiated superficial cells ^14,15^. E1 and E2 proteins are responsible for viral DNA replication and partitioning of viral genomes in host cells, mainly the basal layer ^11,16–20^. E5, E6, and E7 proteins are required to evade the innate immune responses of the host cells and to create an environment to support HPV replication in the superficial layers ^11,17–21^. These genes and their induced cellular changes with cell proliferation, apoptosis, genomic stability, and transformation directly or indirectly contribute to host cell transformation, cancer initiation, and malignant phenotype maintenance ^13,22–36^. To date, many processes that underlie the HPV life cycle and oncogenesis remain largely unknown due to a lack of systems that model natural HPV productive and abortive life cycles. There are several cancer lines with HPV16 or 18, keratinocyte systems (immortalized HaCat cells or primary keratinocytes) with transfection of HPV genomes, keratinocytes (immortalized HaCat cells or primary keratinocytes) infected with quasivirus, pseudovirus, or “native” HPV particles, or immortalized cell lines with E6/E7, SV40, hTERT, while there are only two or three patient-derived cell systems, for example, W12 (HPV16) and 9E (HPV31). The precise process of HPV infection and oncogenesis can be experimentally determined and validated on patient-derived clinically relevant models, thus, making such a model is critically needed for studies of HPV biology and oncogenesis. Our group and others ^37–42^ recently established a clinically relevant cellular model with patient-derived, naturally infected HPVs, 3D ALI cultures from these PCDs employ the HPV life cycle and have been used for anti-HPV screening ^43,44^. Long-term HPV/host interactions, especially proliferation manipulated by the interactions, become at high risk for the transformation of the host cells, which is not a primary intent of HPV infection. E6 and E7 proteins from high-risk HPVs are essential for immortalizing keratinocytes and the progression to transformation ^35,36^. The two oncogenes are expressed in cervical cancer cell lines and primary and metastatic cervical cancers ^45–47^. A deep understanding of the etiology of HPV infection in cervical cancer is critical to the prevention and treatment of the disease.

Single-cell RNA-sequencing (scRNA-seq) has been widely applied to cancer research, including cervical cancer ^48–50^, with incomparable advantages in revealing cellular heterogeneity. The key benefit of using single-cell sequencing for microbial profiling lies in the capability of cell barcodes to match microbes with their respective somatic cells ^51^. Several studies have been carried out to investigate HPV infection in cervical cancer by deriving HPV genes from host scRNA-seq data. These studies either investigate the heterogeneity between HPV-positive (HPV+) and negative (HPV-) tumor cells ^50^, or the immune cell responses and microenvironment changes in HPV+ cervical cancer ^52^ and head and neck squamous carcinoma ^53^. However, there is still a lack of systematic investigations of host cell differentiation and viral gene-specific heterogeneity in cervical cancer. Several questions remain unsolved: Are the expression of HPV genes tending to be epithelial differentiation marker-specific? Is there any epithelial heterogeneity associated with HPV gene expression? How individual viral gene expression affects host gene expression profiling in different cell clusters?

To answer the above questions, we investigate the epithelial heterogeneity and predict epithelial-specific gene signatures in an HPV+ host scRNA-seq data from a human cervical cancer sample. Our result shows evidence of novel cell layers in HPV+ cervical cancer with dysregulation of cell differentiation, extracellular matrix (ECM) structure, and cell cycle. These undefined epithelial layers maintain a high level of E6 and E7 gene expressions and abnormal expression levels of normal layer signature genes. We also explored the heterogeneity between HPV+ and HPV-cells in each epithelial cluster and found HPV+ cells favor epithelial-mesenchymal transition, extra-epithelial matrix, and cellular differentiation signalings in one specific epithelial layer. Our study presented a novel perspective on the investigation of HPV gene expressions in cervical cancer and their interaction with epithelial layers.

## Materials and Methods

### Collection and processing of the scRNA-seq dataset

The scRNA-seq data was collected and sequenced from a 53-year-old female patient who had undergone resection in the Hospital of Fudan University, China. The raw sequence fastq files were downloaded from GSE168652 in the Gene Expression Omnibus (GEO) database ^17^. The data contained 11,289 cells derived from one sample of cervical cancer tissue. The raw sequencing FASTQ files were aligned to the GRH38 reference genome using Cell Ranger ^54^ (10X Genomics, version 7.1) to produce a host gene expression matrix by the STAR algorithm ^55^ (**Fig. 1**).

**Fig. 1.**
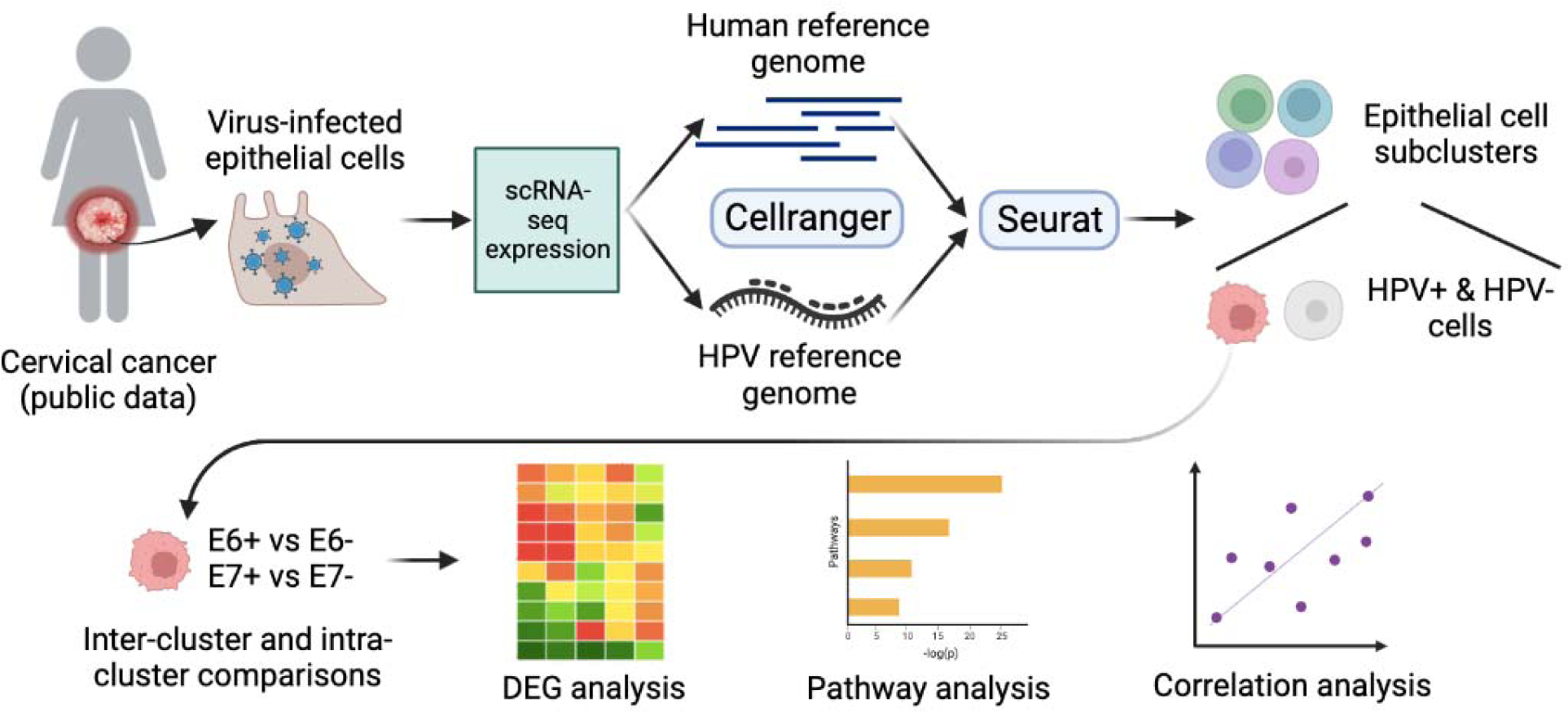
Workflow diagram of the study. Host scRNA-seq data acquired through the public domain from one sample of cervical cancer tissue were aligned to human and HPV reference genomes, respectively. The sequencing expression matrix of the host and virus was generated and combined into Seurat for cell clustering and annotation. Inter-cluster and intra-cluster comparisons were analyzed on epithelial cell clusters based on whether the cells were HPV+ or HPV-. The HPV+ epithelial cells were further divided into E6+/E7+ and E6-/E7-cells. Downstream analyses such as differential expression analysis, pathway enrichment analysis, and gene expression correlation analysis were conducted to reveal the cellular heterogeneity of HPV-infected epithelial cells and the relationship between host and viral gene expressions.

### Sequence alignment of HPV

The complete viral genome and annotation files were downloaded from the NCBI database. The reference sequences for HPV type 16 (HPV16; NCBI, NC_001526.4, 7906bp) and type 18 (HPV18) (HPV18; NCBI, NC_001357.1, 7857bp) were used as the viral reference genomes for alignment. The “mkref” function in Cell Ranger (10X Genomics, version 7.1) was used to build the HPV reference files. To detect HPV present in the single cells, we aligned the data to the HPV reference files using the “count” function in Cell Ranger. The parameters “alignIntronMax” and “genomeSAinndexNbases” for the STAR algorithm (integrated into Cell Ranger) were adjusted based on the characteristics of the viral genome. The “alignIntronMax” parameter, representing the maximum intro length allowed in the algorithm, was set to 1 because viruses do not contain introns. To achieve a balance between memory size and speed, the “genomeSAinndexNbases” parameter was set to 6 based on the algorithm min(14, log2(genome length)/2-1). The “genomeSAinndexNbases” was set to 6 bases on algorithm min(14, log2(genome length)/2-1) to balance the computational memory size and speed. Subsequently, the count matrices of both host and virus genes were integrated using a single-cell index for downstream analyses (**Fig. 1**).

### Preprocessing and cell clustering of scRNA-seq data

The raw count matrix was processed using Seurat (version 4.3.0) package ^56^ in R (version 4.2.1). Transcripts were detected in at least three cells, and cells containing at least 200 transcripts were processed. The cells that had over 6,000 expressed genes and the genes that over 25% of UMIs mapped to the mitochondrial genome were removed. Next, the gene expression matrix was log-normalized, and the top 2,000 highly variable genes were selected as features for principal component (PC) analysis. Cells were then clustered using the Louvain clustering method on the top 20 PC dimensions with a resolution of 0.6, and UMAP non-linear dimensional reduction was performed for visualization. Cells that have at least one of the HPV genes expressed were labeled as HPV+ cells, and cells that contain only E6 or E7 HPV transcript were labeled as E6+ or E7+ cells.

### Differentially expressed genes analysis

Differentially expressed genes (DEGs) analysis among clusters in subsets of all the cells was performed using the FindAllMarker function in the Seurat package based on the Wilcoxon Rank Sum test. DE analysis between cellular groups was performed using the FindMarker function. The log2 fold change is calculated with the absolute threshold of 0.25. Significant DE genes were defined with Bonferroni-adjusted *p*-values <0.05.

### Functional enrichment analysis

DEGs were loaded into enrichR ^57^ for the Gene Ontology (GO) enrichment and Kyoto Encyclopedia of Genes and Genomes (KEGG) pathway analysis. GO terms for biological progress (BP) and molecular function (MF) gene sets were based on Gene Ontology (GO) knowledgebase. Any terms with an adjusted *p*-value lower than 0.05 were reported as significantly enriched pathways.

### Gene expression correlation analysis

The relationship between host and viral genes, specifically E6 and E7, was investigated using Pearson’s Correlation analysis. Host genes selected for this study exhibited the most significant differential up-regulation within each epithelial cluster compared to the remaining ones. P-values were adjusted using the Benjamini-Hochberg approach. The criteria for determining significant correlations were an absolute coefficient value exceeding 0.25 and an adjusted *p*-value less than 0.05.

## Results

### HPV genes differentially expressed in epithelial cell clusters

A total of 11,399 cells were clustered into 11 groups (**Fig. 2A**). We manually annotated these cell clusters based on public signature genes, including nine epithelial cell clusters (*CDH1*, *EPCAM*, *CDKN21*, *KRT8*), one CD8+ T cell cluster (*CD3D*, *CD3E*, *CD3G*, *CD8A*, and *NKG7*), and one macrophage cell cluster (*CD163*, *CD14*, *MRC1*) (**Fig. 2B**). As described in the original paper, the scRNA-seq data was obtained from the tumor region of a patient with cervical cancer; thus most cells expressed markers of epithelial cells^48^. The CD8+ T cells and macrophage cells may be residues that failed to be isolated from the tumor cells. HPV16 transcripts (E1, E6, E7) were detected through mapping to the reference genome, while HPV18 genes were not detected. HPV16 was predominantly associated with the epithelial cell clusters. In particular, Epi_2, Epi_5, and Epi_8 had the highest proportion of HPV+ cells (33.6%, 37.5%, and 32.4%, respectively) (**Fig. 2C and Supplementary Table S1**). Epi_7 and Epi_10 exhibited the lowest proportion of HPV+ cells (7.21% and 5.63%, respectively). The identified HPV16 transcripts were enriched in E1, E6, and E7 genes, consistent with the findings of another study utilizing the same dataset ^58^. There were significant levels of HPV gene expression in epithelial cells than in CD8+ T cells and macrophages (**Fig. 2D-G**), suggesting heterogeneity of HPV gene expression in HPV+ cancer cells.

**Fig. 2.**
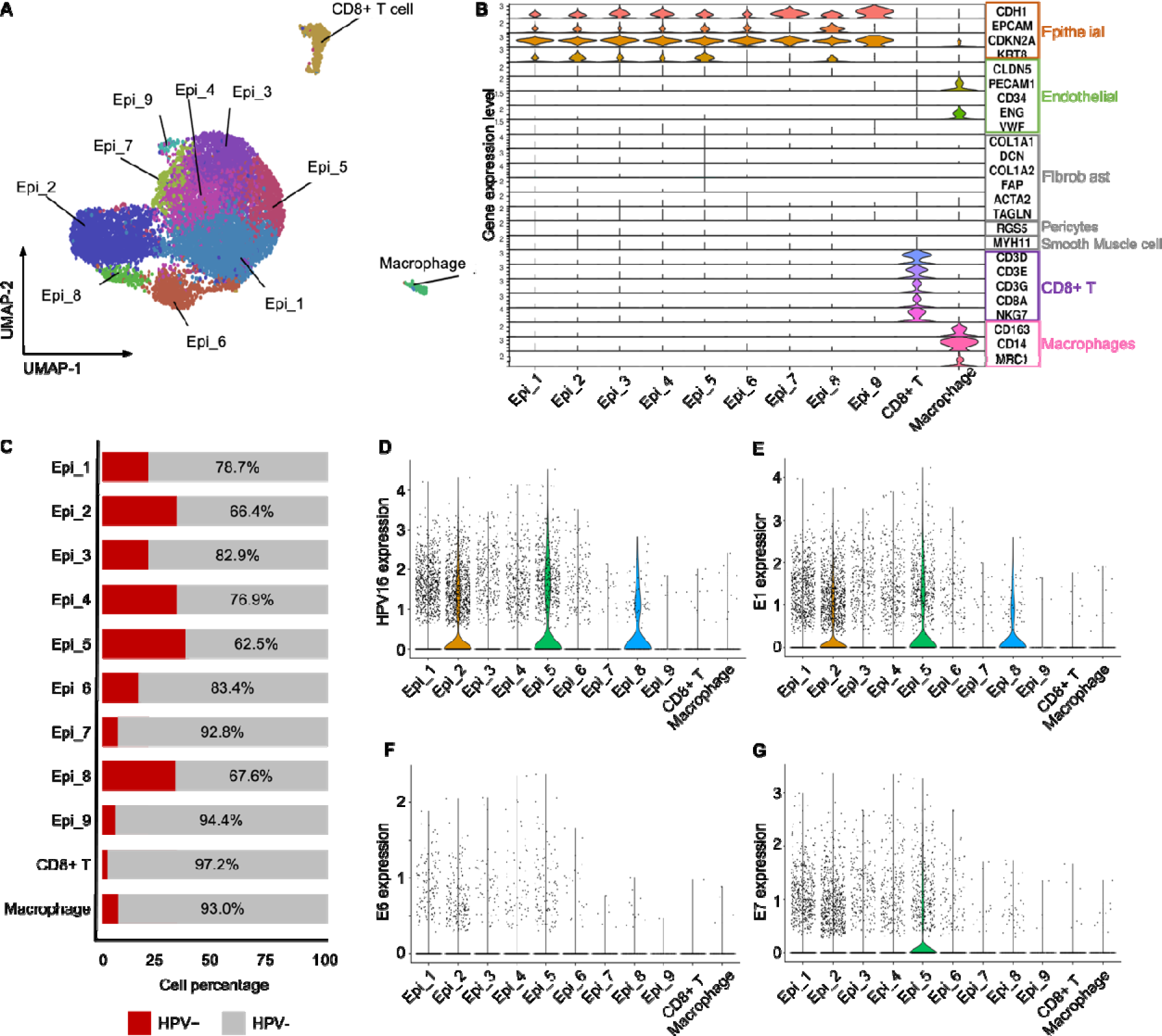
Differential HPV gene expression in epithelial cell differentiation-related clusters. **A)** UMAP plot of host cell clusters and annotation; **B)** The Violin plot of typical cell-type markers across all clusters; **C)** The bar graph quantifies the proportion of HPV+ and HPV-cells in cell clusters; **D)** Violin plot indicating the expression of viral transcriptomics; **E**) Violin plot indicating the expression of E1 gene expression; **F**) Violin plot indicating the expression of E6 gene expression; **G**) Violin plot indicating the expression of E7 gene expression.

### Intra-tumoral heterogeneity of epithelial cells with HPV infection

To characterize the effects of HPV transcripts displayed in the epithelium, we included epithelium clusters (Epi_1-9) with 2,801 cells in the subsequent analysis. As specific cervical epithelium markers have not been reported, we used gene makers of normal human skin epithelium^59^ to further distinguish the nine epithelial cell clusters, including basal layer (*MKI67*, *KRT5*, and *TP63*), spinous layer (*TGM1* and *TGM5*), granular layer (*FLG*, *KRT1*, *KRT2,* and *KRT10*), and corneum layer (*IVL*, *PPL,* and *EVPL*) (**Fig. 3A**). As a result, Epi_2 and Epi_8 displayed basal-like attributes with specific expression of *MKI67*. The rest epithelial clusters could not be classified into a specific cellular layer as the expression pattern of these markers was not clear. Epi_3 and Epi_4 are spinous-like as they expressed markers *TGM1* and *TGM5* of spinous layers. Epi_1 and Epi_6 are granular-like. However, Epi_1, Epi_3, and Epi_4 also express gene *KRT1*, a marker for the granular layer. We also noticed that neither of these markers was expressed in Epi_5, the cluster with the highest proportion of HPV+ cells and elevated viral gene expression levels. On the contrary, Epi_7, a cluster characterized by a low HPV+ cell ratio and diminished viral gene expression, showed the expression of almost all epithelial markers. It is intriguing to note the almost complete absence of epithelial layer markers in Epi_5, while they are manifest in Epi_7. Additionally, Epi_9 showed similar expression levels as Epi_7 in the stratum corneum layer. The epithelium cells were not ordered as in normal epithelium layers, indicating potential aberrations in cell arrangement due to HPV/host cell interactions during cell differentiation, transformation, and cancer development. Our observations indicated that significant heterogeneity exists among the cells in cancer tissues and that these cells potentially exhibit unique/different gene signatures compared with normal epithelial cells.

**Fig. 3.**
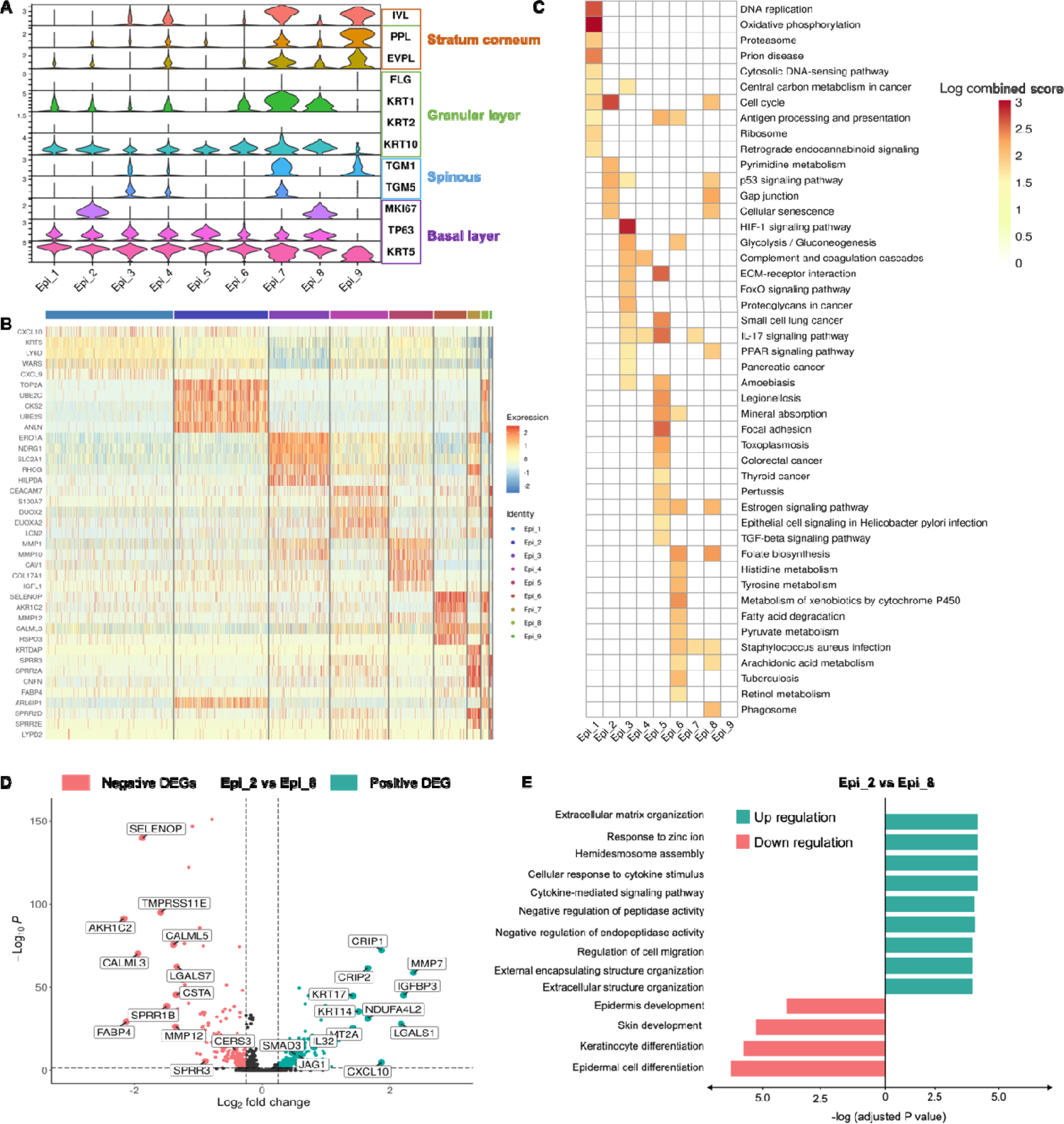
Identification of intra-tumoral heterogeneity. **A)** Violin plot of collected cell-layer marker genes; **B)** The expression levels of the top 5 up-regulated DEGs in each epithelial cluster; **C)** Functional enrichment analysis for KEGG showing the up-regulated signaling pathways differential regulated across epithelial clusters, the color scale indicating log combined scores of the pathway, value zero indicating no pathway identified in the corresponding cluster; **D)** Volcano plot showing the differentially expressed genes between Epi_2 over Epi_8. The dashed line indicates the threshold of significant gene expression, defined as the absolute value of log fold change >0.25 with -log10(*p*) >1.3; **E)** BP in GO term showing the signaling pathway that is differentially regulated in cells of Epi_2 versus Epi_8.

The identified up-regulated DEGs between epithelial clusters were used to characterize expression pattern crossing epithelium (**Fig. 3B and Supplementary Table S2**). Heterogeneity in epithelial cell gene expression was detected by gene set enrichment analysis of the top 100 DEGs defining each cluster (**Fig. 3C**). Both Epi_2 and Epi_8 showed significant enrichment of the p53 signaling pathway, gap junction, and cellular senescence. Epi_8 additionally enriched in pathways associated with estrogen signaling and staphylococcus aureus infection. Furthermore, cells from Epi_4 exhibited significant upregulation signaling pathways that are involved in response to microbial infection, such as amoebiasis, legionellosis, and pathways related to cancer and estrogen signaling, as well as cell adhesion via the ECM receptor interaction and focal adhesion. Estrogen signaling and ECM production play a significant role in the development and progression of cervical cancer ^60^. In general, the epithelial clusters exhibited heterogeneity in signaling functions. Specifically, Epi_5 has the highest ratio of HPV+ cells and shows high associations with HPV oncogene expression, suggesting a highly malignant epithelial cell population. Epi_2 and Epi_8 both exhibited basal-like attributes but slipt into different epithelial clusters. To find out how the two slipt epithelial clusters affect epithelial signaling pathways, cells in Epi_2 were compared to cells in Epi_8. We showed that the Epi_2 was characterized by upregulation of the extracellular structure molecule *MMP*, the cell migration gene *JAG1*, and chemokines *CXCL10* and *CCL20* (**Fig. 3D and Supplementary Table S3**). In addition, we observed a significant downregulation of epidermis cell and keratinocyte differentiation, and epidermis development in GO enrichment analysis, with decreased expression of molecules from small proline-rich proteins family (**Fig. 3E**). A comparison between basal-like epithelial clusters showed that gene expression and cell signaling pathways related to epithelium structure were modestly affected.

### Differential expression of host genes in epithelial clusters associated with E6 and E7 genes

Even though Epi_5 has the highest ratio of HPV-infected cells compared with other epithelial clusters, any epithelial-layer markers could not define Epi_5, and the cluster’s layer characteristics are unclear. To investigate host transcriptomics in the HPV+ cells, we subset HPV+ cells and compared these cells in Epi_5 with the remaining epithelial clusters. HPV+ cells in Epi_5 expressed high levels of ECM-associated genes (*MMPs*, *COL17A1*, and *ITGB1*), cell adhesion genes (*LAMC2* and *BCAM*), cell proliferation genes (*F3*), and apoptosis-related genes (*G0S2* and *TNFRSF12A*), which are related to aberrant epithelial cells growth and adhesion (**Fig. 4A and Supplementary Table S4**). MMP10 can regulate tumor cell migration and invasions which may cause resistance to apoptosis via apoptotic pathways in cervical tumors ^61^. *COL17A1* encodes an essential component in the ECM called collagen XVII. A study reported collagen XVII overexpression in cervical tumor ^62^. E7 displayed a similar DEG expression pattern to HPV+ cells within the Epi_5 (**Fig. 4A and Supplementary Table S4**). The ontological enrichment analysis of the up-regulated genes revealed that HPV+ cells in Epi_5 were highly enriched in processes related to ECM organization and disassembly, cell cycle regulation, epithelial cell proliferation, and the regulation of extrinsic apoptotic signaling pathway (**Fig. 4B**). Interestingly, only a few DEGs (including *COL17A1*, *BCAM*, *RND3*, *PSG5*, *CAV1*, and *PPP1R15A*) were identified in the E6+ cell population (**Fig. 4A**). This suggests that E7 is the principal viral gene contributing to the observed characteristics in the epithelial tumor tissue, confirming a previous study ^22^. Therefore, the Epi_5 cluster showed noticeable dysregulation in ECM structure, cell cycle, and apoptosis, possibly attributable to HPV infection. These alterations in cellular functionality potentially facilitated tumor invasion within Epi_5. This may explain why normal epithelial markers were absent in this cluster, indicating a significant change in cell behavior and the heterogenic malignancy of this cell population.

**Fig. 4.**
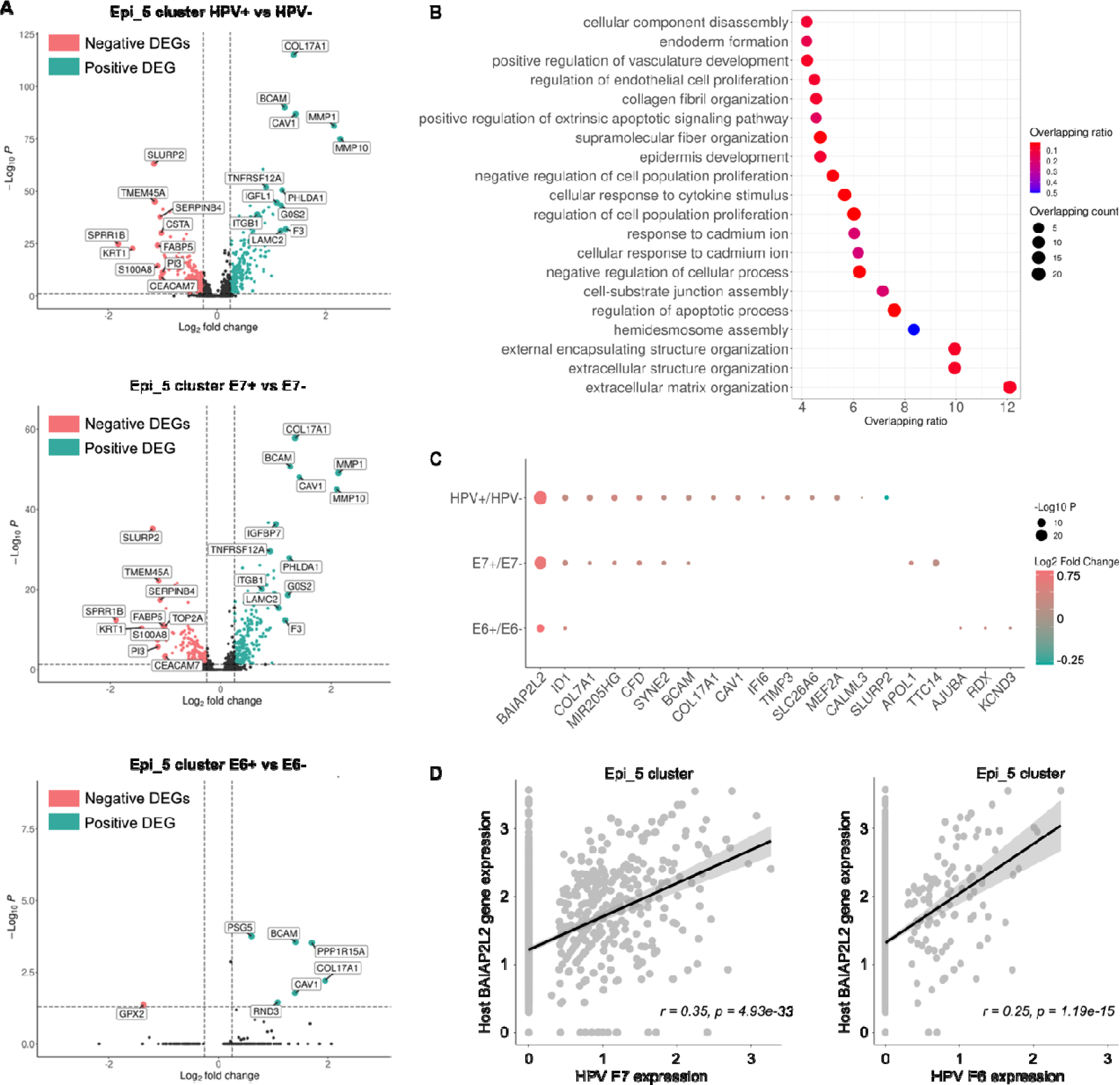
Epi_5 regulated by HPV+ cells. **A)** Differentially expressed genes of Epi_5 over other clusters in HPV+, E6+, and E7+ cells, respectively; **B)** GO enrichment pathway of biological processes of up-regulated DEGs in HPV+ cells over the rest clusters. **C)** DEGs of HPV+ over HPV-cells, E7+ over E7-cells, and E6+ over E6-cells, respectively. **D)** Host genes’ correlation regression with E7 and E6 expression in Epi_4, respectively; r: correlation coefficient; *p*: adjusted *p*-value.

To explore the HPV oncogene-related heterogeneity in Epi_5, HPV+ epithelial cells were directly compared to HPV-epithelial cells. We found that Epi_5 cells with positive HPV transcripts demonstrated a significant elevation in the expression of genes involved in the cytoskeleton organization (*BAIAP2L2*), epithelial-mesenchymal transition (EMT) regulation (*MIR205HG*, *TIMP3*, and *BCAM*), ECM structure (*COL7A1*), and ion channel modification (*CAV1*) (**Fig. 4C and Supplementary Table S5**). *BAIAP2L2* was also up-regulated in Epi_1, Epi_2, Epi_3, Epi_4 (**Supplementary Table S5**). This gene was found to be active in cervical squamous cell carcinoma, with high-level expression indicative of unfavorable overall survival in patients with this carcinoma ^63^. Although there is no direct association linking *CAV1* with cervical cancer, DNA viruses, including HPV16, John Cunningham virus, and bovine papillomavirus type 1 (BPV1), have been found to enter host cells through clathrin-mediated endocytosis and CAV1-1-dependent pathways ^64^. In addition, *MIR205HG*, which regulates EMT, may influence epithelial cells in cervical cancer. Collectively, these results indicate HPV16 predominately exerts its influence on the Epi_5 by modulating gene expression associated with cell differentiation and organization. The reaming epithelial clusters displayed functional differences between HPV+ and HPV-cells (**Supplementary Table S5**).

Understanding how the HPV gene interacts with host genes is also essential. The comparisons were performed on the subsets of E6+ or E7+ epithelial cells in Epi_5. The *ID1* gene, which was found to be up-regulated in HPV+ cells, showed significant elevation in both E7+ and E6+ cells (**Fig. 4C and Supplementary Tables S6-7**). *ID1,* associated with cell differentiation and DNA-binding activity, has been noted up-regulated in cervical cells ^65^. Overexpression of *ID1* has been linked to HPV E6 in invasive breast cancer patients ^66^. Moreover, ID1 has been suggested as a potential oncogene due to its increased expression in cervical cancer ^67^. Another gene, *BAIAP2L2,* showed pronounced upregulation in the three HPV+ groups, suggesting that *BAIAP2L2* may play a distinctive role in HPV+ cervical cancer. *MIR205HG* showed up-regulated in E7+ cells, but not in E6+ cells, indicating that E7 is the primary contributor to the miR-205-5p associated pathway to regulate EMT. E7+ cells also demonstrated increased expression of genes *BCAM* and *COL7A1*, which enriched in both EMT and ECM regulation, suggesting the viral E7 has modified Epi_5 by changing epithelial structure and altering ECM (**Fig. 4C and Supplementary Table S6**). Additionally, gene *IFI6* was overexpressed in HPV+ cells comparing HPV-cells (**Fig. 4C**), but was not identified in E7+ or E6+ cells. However, the expression of *IFI6* was observed to be elevated in cells transfected with HPV16 E7, and a correlation was found between the high expression of this gene and reduced patient survival ^68^. This suggests that HPV infection may alter the host gene IFI6 via additional viral genes apart from E7. *IFI6* is involved with the ERBB signaling pathway, enriched in HPV+ cells. The critical protein ErbB2 has been identified within this pathway as a negative prognostic factor in human cervical cancer^69^. Therefore, *IFI6* may be linked with ErbB2, and the protein it directly encodes could potentially be targeted for cervical cancer treatment. In conclusion, E6 and E7 modify epithelial function by impacting different pathways and collectively contribute to cervical cancer progression. Beyond the roles of E6 and E7 genes, HPV interacts with cervical cancer host epithelial cells regarding host entry and ERBB signaling.

We found that epithelial clusters displayed heterogeneity in HPV-associated cervical cancer; specifically, Epi_2, Epi_5, and Epi_8 displayed altered structures. To further investigate if viral genes were associated with host genes, we performed a comprehensive correlation analysis comparing the expression of viral genes (E6 and E7) with significant up-regulated DEGs from each epithelial cluster. Our data reveal that gene BAIAP2L2 is related to both E6 and E7 gene expression in Epi_5 (r=0.25, r= 0.35, respectively) (**Fig. 4D**). It corresponded to the above finding that *BAIAP2L2* was up-regulated in HPV+ cells in Epi_5. In Epi_8, gene *CKAP2L* (r=0.29, adjusted p=0.038) showed a correlation with E7 expression (**Supplementary Table S8**). However, no genes were identified as significantly correlated with E6 or E7 expression in Epi_2 and other epithelial clusters.

## Discussion

In this study, we investigated the heterogeneity of epithelial cells and gene signatures specific to these cells in a sample of HPV-infected cervical cancer with scRNA-seq, facilitating the understanding of the interaction between HPV and cervical epithelium differentiation. Our investigation not only involved a comparison between HPV+ and HPV-cancer cells but also observed the effects of the individual viral genes, E6 and E7, in cancer cells. Additionally, we estimated HPV16 gene expression in cervical cancer and identified specific epithelial cell clusters - specifically Epi_2, Epi_5, and Epi_8 - predominantly positive with HPV transcripts. Since we did not examine the status of HPV genomes in these cells, we were not able to rule out whether the heterogeneity of HPV gene expression was due to the sensitivity of RNA-seq or natural expression differentiations. In some cases, the expression of viral E6 and E7 may be under detectable levels, due to alternative splicing. Subsequently, we delved into the epithelial heterogeneity among these clusters. Epi_2 and Epi_8 exhibited characteristics of basal-like epithelium, while Epi_5 did not align with any typical epithelial layer markers.

Given that HPV initiates its activity in the basal layer of the cervical epithelium upon entering host cells, it is plausible that Epi_2 and Epi_8 reflect an enrichment pathway triggered by HPV transcripts, including the p53 signaling pathway. These two basal-like clusters display differences in gene signatures related to ECM structure and keratinocyte differentiation, suggesting altered epithelial structure by HPV transcripts. When looking at the co-expression relationship of the host gene and individual viral gene, we only found host gene *CKAP2L* was positively correlated with E7. *CKAP2L* was found to be highly expressed in hepatocellular cancer, lung adenocarcinoma, and breast cancer ^70–72^, where *CKAP2L* was proposed as a potential target gene for tumor therapy. We hypothesized that *CKAP2L* might also participate in the development of cervical cancer. The epithelial structure of the cervix contributes to its vulnerability to infection, and distinct epithelial cells in different epithelial sites may regulate viral gene expression differently^73^. Interactive manipulation of specific HPV genes and host gene transcriptomics can alter normal epithelial function, highlighting the complex interplay mechanisms behind the virus and host.

We paid particular attention to Epi_5 due to its highest proportion of HPV+ cells and the lack of alignment with normal epithelial layer markers. In Epi_5, high HPV expression led to changes in the expression of genes related to ECM, cell adhesion, proliferation, and apoptosis, including *BAIAP2L2, MMPs, COL17A1, ITGB1, F3, G0S2, and TNFRSF12A*. Interestingly, many of these genes have been linked to the progression and severity of various types of cancers, including cervical cancer. As E7+ cells showed similar expression patterns in HPV+ cells, indicating viral gene E7’s key contribution to epithelial tissues in cervical cancer. Through analyzing the correlation between individual host genes and viral genes, we found that *BAIAP2L2* exhibited a strong association with E7 and was highly regulated in HPV+ cells or cells expressing individual viral genes. This suggests that the biological pathways and cascade reactions induced by *BAIAP2L2* warrant further exploration in experimental studies. Overall, our study provides a unique viewpoint on examining epithelial heterogeneity and the role of HPV16 in cervical cancer. We uncovered differential HPV expression across epithelial clusters, demonstrating that HPV-infected epithelial cells exhibit dysregulation in cell differentiation and structural organization. The host gene expression profiling varies across epithelial differentiation layers in response to individual viral genes. Computational identifications in this manuscript can guide the design of naturally HPV-infected models, enabling the identification of interactions between intratumoral virus and pathological associated immunity and metabolism.

Our study still has room for improvement regarding the analysis and characterization of layer-specific epithelial markers. First, our research relied on a single dataset; therefore, the findings need validation in additional cohorts to deepen our understanding of HPV and cervical cancer. As more datasets become available, we intend to corroborate our inferences further. Second, Viral gene E1 is critical for the HPV life cycle. In our future study, the roles of E1 in viral persistence, cancer initiation, and development could be studied with such models. Third, while our study identified correlations between HPV E6/E7 genes and host gene expression, the underlying mechanisms are yet to be elucidated. This could be achieved by integrating dynamic viral transcriptomics cases or establishing experimental cell models to mimic host cell differentiation. It might also be insightful to investigate the relationship between the genomic insertion position of viral genes and host gene promotor/enhancer regions. Fourth, more comprehensive mechanistic studies are required to fully understand the intricate interactions between HPV genes and host gene expression during persistent viral infection and transformation. The development of spatial transcriptomics and the application of cutting-edge deep learning models^74^ present an exciting opportunity in analyzing host-resolved viral infection. Lastly, precise processes of HPV life cycle and oncogenesis still remain largely unknown due to the lack of natural systems for HPV productive and abortive life cycles. It would be important to generate more patient-derived clinically relevant cell models ^38,42,75,76^.

## Author Contributions

Drs. Qin Ma and Xuefeng Liu conceived the idea. Yingjie Li performed data analysis and wrote the manuscript draft. Cankun Wang provides support for coding. All authors revised and edited the manuscript, and approved the final version of the paper.

## Funding Statement

This study was supported by the National Institutes of Health R01GM131399 (Q.M.) and R01CA276474, R01CA222148, R33CA258016 (X.L.). This work was supported by the Pelotonia Institute of Immuno-Oncology (PIIO). The content is solely the responsibility of the authors and does not necessarily represent the official views of the PIIO.

## Data Availability Statement

The dataset used in this paper can be found in the GEO database (GSE168652).

## Conflict of Interest Disclosure

The authors declare no conflict of interest.

## Supplemental Tables

Supplementary Table S1: cell counts and percentage of HPV-positive cells crossing clusters.

Supplementary Table S2: Differentially expressed genes in each epithelial cluster.

Supplementary Table S3: Differentially expressed genes between Epi_1 and Epi_8.

Supplementary Table S4: Differentially expressed genes in Epi_5 of HPV+, E7+, E6+ cells.

Supplementary Table S5: Differentially expressed genes between HPV+ and HPV-cells in Epi_5.

Supplementary Table S6: Differentially expressed genes between E7+ and E7-cells in Epi_5.

Supplementary Table S7: Differentially expressed genes between E6+ and E6-cells in Epi_5.

Supplementary Table S8: Pearson’s correlation analysis of selected host genes with viral E6 and E7, respectively.

## Notes

### Competing Interest Statement

The authors have declared no competing interest.

