## Supplementary Table S1 for "Identification of HPV oncogene and host cell differentiation associated cellular heterogeneity in cervical cancer via single-cell transcriptomic analysis"

| **Clusters** | **total cells** | **Pos** | **% Pos** | **E1** | **% E1** | **E6** | **% E6** | **E7** | **% E7** |
| --- | --- | --- | --- | --- | --- | --- | --- | --- | --- |
| Epi_1 | 3320 | 707 | 21.3 | 650 | 19.58 | 133 | 4.01 | 477 | 14.37 |
| Epi_2 | 2439 | 819 | 33.58 | 779 | 31.94 | 178 | 7.30 | 581 | 23.82 |
| Epi_3 | 1563 | 267 | 17.08 | 251 | 16.06 | 62 | 3.97 | 206 | 13.18 |
| Epi_4 | 1501 | 347 | 23.12 | 322 | 21.45 | 66 | 4.40 | 233 | 15.52 |
| Epi_5 | 1154 | 433 | 37.52 | 417 | 36.14 | 107 | 9.27 | 322 | 27.90 |
| Epi_6 | 844 | 140 | 16.59 | 134 | 15.88 | 22 | 2.61 | 95 | 11.26 |
| Epi_7 | 319 | 23 | 7.21 | 22 | 6.90 | 6 | 1.88 | 16 | 5.02 |
| Epi_8 | 188 | 61 | 32.45 | 57 | 30.32 | 11 | 5.85 | 41 | 21.81 |
| Epi_9 | 71 | 4 | 5.63 | 3 | 4.23 | 1 | 1.41 | 3 | 4.23 |
| CD+ 8 T cells | 392 | 11 | 2.81 | 10 | 2.55 | 1 | 0.26 | 6 | 1.53 |
| Macrophages | 142 | 10 | 7.04 | 9 | 6.34 | 4 | 2.82 | 8 | 5.63 |

Table.S1. **cell counts and percentage of HPV-positive cells crossing clusters.**
